# ISiCell: involving biologists in the design process of agent-based models in cell biology

**DOI:** 10.1101/2023.06.30.547165

**Authors:** Florian Cogoni, David Bernard, Roxana Kazhen, Salvatore Valitutti, Valérie Lobjois, Sylvain Cussat-Blanc

**Author notes:** These authors contributed equally to this work (Florian Cogoni, David Bernard).

## Abstract

Agent-based models are commonly used in biology to study tissue-scale phenomena by reproducing the individual behavior of the cells. They offer the possibility to study cellular biology at the individual cell scale to explore the basic behavior of cells which are responsible of the emergence of more complex phenomena at the tissue scale. Additionally, they can produce a predictive tool that will help taking decisions for biologic experiments based on *in silico* simulations. However these models require a good intercomprehension between the biologists and the modelers and thus it may take weeks or months to end up providing a usable prototype.

To address this limitation, we propose a new methodology to facilitate the dialog between biologists and modelers and improve biologists’ involvement in the design of the model. For this purpose, UML diagrams, in particular, state-transition and activity diagrams, are used. They allow a better comprehension of the model for the biologists and offer a general frame for structuring models. Visualization of simulations is also used to have qualitative feedbacks from the biologist on the model. They are instrumental to validate or refine the prototype before exploring it.

Alongside this methodology, we propose a web platform that enables to build state-transition and activity diagrams to describe a model and translate them into code. The generated code is then compiled on-the-fly and simulations are ready to visualize and explore. The platform also disposes of tools to directly visualize and manually explore the model. These tools allow for qualitative validation of the model and additional interaction with the biologists.

Finally in this article, we show the capacity of our platform to reproduce models from the literature and to build new models starting from workshops with biologists. Its range of application is wide and includes immunology, oncology or cell biology.

**Author summary:** We developed a methodology based on diagrams to facilitate the dialog between computer scientists and biologists when building *in silico* models. The main idea is to limit misunderstandings and improve the involvement of the biologists in the prototyping process. For this purpose, we use visual methods to simplify the modeling phase. Alongside this methodology, we propose a web platform, called ISiCell, which enables to visually code thanks to diagrams that will be translated into code. The platform allows for compiling the generated code on the fly and to visualize and explore the model directly with the platform. The strong advantage of the platform is that one day workshop biologist/modeler allows to build new models. Additionally, we were able to reproduce models from the literature within the modeling platform showing the versatility of the tool.

Our long-term objective is to use our methodology and platform in new contexts to develop new models. We intend the make the platform more user friendly in order to expand the community of users. Involving biologists in the conception of *in silico* models might improve their acceptability in the community.

## Introduction

### Interactions between computer scientist and biologist

Models have long-standing history in helping domain experts to better understand and predict phenomena in complex systems. In biology, they allow to highlight physiological mechanisms in living organisms. Significant progress has been possible by the use of animal models (*in vivo* model) allowing to represent a realistic physio-pathological context. However, these experiments lead to significant economic and ethical costs, and are subject to inter-individual variability which may require many replicates. To overcome these constraints, biologists also use cell cultures (*in vitro* model) which allow lower costs and better control of experimental conditions, while limiting inter-experimental variability. In recent years, mathematical and now computational tools have allowed a new abstraction: *in silico* modeling. Although these models are still difficult for biologists to use and apprehend, they allow the exploration of working hypotheses with relatively low costs and precise control of experimental conditions. These models make it possible to aggregate different kind of experimental data while formulating and testing biological hypotheses. They serve two purposes. Firstly, they reinforce the pre-existing knowledge by challenging previously assumed hypotheses by matching biological data with a calibrated computational model. Secondly, they serve as a predicting tool allowing to explore and find specific and interesting *in silico* cases before experimentally evaluating these conditions. All these models are not mutually exclusive and should be, on the contrary, complementary.

A good understanding of biological functions is achieved if all relevant information at multiple levels of organization is integrated, as described in [1–3]. The need to model complex spatio-temporal processes at many scales has led to the emergence of many computational modeling techniques, including systems of differential equations, cellular automata simulators and Agent-Based Models (ABM) [4]. While, differential equations allow for a mathematical description of the model at a population scale (but fail to describe individual-level phenomena) cellular automata allow to discretize the environment allowing to spatially study local and individual mechanisms. ABM have many advantages over those methods for modeling cell biology phenomena, especially for testing hypotheses at the cell scale. We will present these advantages in the following subsection.

### ABM in biology

Agent-based modeling is a widely used approach to quantitatively simulate dynamic systems [5]. Over the last decade, this modeling approach has become more popular and has been applied to a wide range of biological systems [6–8]. Each ABM is defined by a set of autonomous agents and a number of stochastic or deterministic rules. These rules govern the interactions of each agent with their neighbors and their environment. Unlike equation-based approaches, ABMs are decentralized, meaning that the overall behavior of the system emerges from the collective behavior of each individual agent in the system.

Each cellular agent can follow a completely independent behavioral trajectory governed by individual parameters reflecting the heterogeneity of a cell population. Modelers can directly implement cellular rules that reflect behavior observations at the cellular level and inter-cellular interactions, allowing them to rapidly translate biological hypotheses into algorithmic rules.

In this way, it is possible to perform simulations that explore the emergent behaviors of these hypotheses and compare them with new data to iteratively confirm, reject or improve the underlying hypotheses [9–11].

Although these models can be non-spatialized, ABMs allow to finely describe the local interactions between cells and the individual influence of these interactions on their behavior. This spatialization enables direct visualization, hence allowing for quick qualitative validation of the produced models.

### Participatory modeling and ISiCell goals

To develop such models, biologists’ knowledge of the cellular scale phenomena must be transmitted to modelers through discussions. However, coding models takes time during which the interactions are nearly impossible, thus inducing iterative production cycles (Fig 1) with biological discussions to guide the modelers followed by a development phase. This differed development can additionally induce misunderstandings translated into faulty behaviors in the model while being difficult for the biologists to raise these conception mistakes. All of these issues combined cause prolonged development phases. This long modeling process may result in reluctance from the biologists to start such a complex and time-consuming collaboration with modelers.

**Fig 1.**
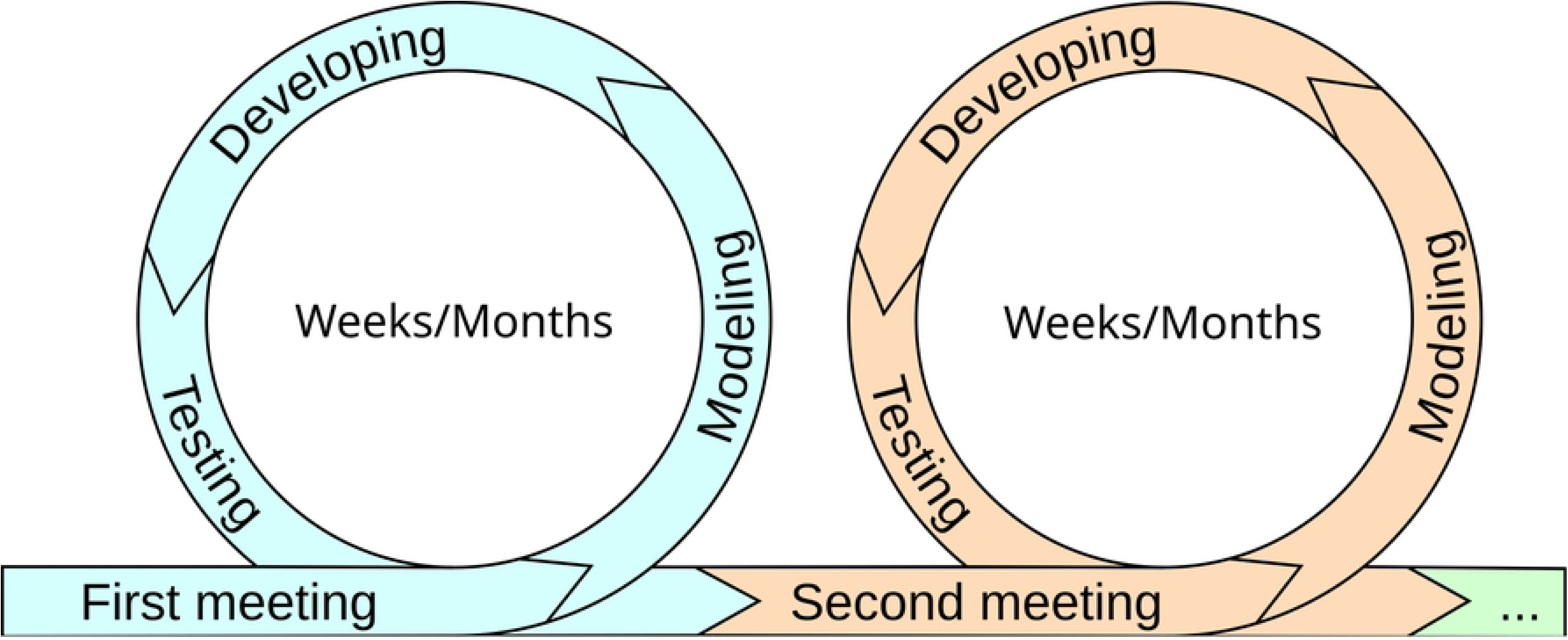
Typical life cycle of the design of biologic models. Each modeling project starts with a first meeting where biologists present their work and their studied phenomena and discuss with the modelers about their needs and expectations from the future model. Then modelers synthesizes what they understood from the biological reality, develop a first prototype and after conceiving a first functional prototype, a second meeting is scheduled to discuss the model and improve it. This process is iterative until abandonment or the obtainment of a satisfying enough model. Each loop can take weeks or months depending on the complexity of the model, misunderstandings and schedules.

Participatory modeling is a paradigm that involves specialist stakeholders in the development process of models by mobilizing their knowledge at each step. This modeling philosophy is commonly used in the fields of environmental resources management [12–14] but, to our knowledge, is currently not applied in the cell biology field. In this approach, ABMs are considered as a tool to achieve a model collaboratively. As a matter of fact, they offer the possibility to easily manage a great number of spatialized, autonomous and decentralized agents which is particularly well suited for socio-ecological systems [15] as it is for modeling biological tissues [16]. In this study, we propose to adapt concepts from participatory modeling to the field of cell biology. To this end, we have developed a methodology that, together with a development platform, allow to co-design with the biologists agent-based model of biological systems at the cell population level.

Our approach was developed to actively integrate the biologists in the prototyping process. This methodology is in line with participatory approaches such as companion modeling [17] and is based on the use of diagrams to facilitate communication and minimize misunderstandings between biologists and modelers.

Alongside the methodology, the ISiCell platform was conceived to support this approach. The main value of ISiCell is to make the biologists proactive actors in the development of the model. This is allowed by its own visual programming paradigm (based on the diagrams of the methodology) facilitating comprehension for biologists without hampering the coding possibilities. The main idea is to confront the biologist with as less code as possible by using visual programming approaches to simplify the intercomprehension and then generate the code on-the-fly from co-constructed diagrams. Subsequently, it also aims at rendering the qualitative evaluation and validation by the biologists of prototypes accessible through simulation visualization. This simulation visualization tool provides a dynamic step-by-step view of the cells and enables to dynamically plot data from the simulation. ISiCell aims at becoming a participatory tool for developing Agent-Based model in the cellular biology field in the same way as CORMAS (COmmon-pool Resources and Multi-Agent Simulations) [18] in the environment and resources management area. The methodology and the platform are evolving in parallel to improve each other.

### Methodology to co-design models

We propose a dedicated methodology with the aim to reduce the design duration of in-silico model. Additionally, we expect our methodology to further implicate biologists experts in the design loop of the models. An additional objective of our methodology is to ensure that the expert biologists fully understand the inner mechanisms of the model by participating to its design process.

This methodology is based on the use of (1) agent-based models, known to be easy to understand for non-expert modelers because based on a set of simple rules at the agent level which produces emerging complex behaviors at the population scale and (2) conceptual schema to graphically represent the behavior of the agents. These are known and proven tools in the domain of participatory modeling to facilitate the communication between modelers and domain experts [19]. Moreover, we propose a web-based modeling platform, ISiCell, that supports this methodology and transform in one-click schema into executable and explorable simulations.

### Graphical representation of the model

Our methodology is based on five successive steps. Even though we suggest to follow these steps in the described order, they can be revised during the modeling session to take into account elements that were not anticipated. Note that the steps are also mostly based on graphical representation to help biologists to apprehend the model.

#### Step 1: environment

In this first step, we determine the environmental conditions of the cells. Modelers and biologists must decide the type of environment to be simulated: 2D/3D, discrete/continuous, physical model (no physics, mass-spring-damper system, Hertzian physics, etc.), molecular diffusion, etc. The time granularity must also be discussed and set up. These choices must be taken into account early on in the design process to evaluate the computational effort necessary to run future simulations.

To guide these choices, modelers need to discuss the maximum number of cells in the simulation, the accuracy required for the physical interactions and any other components that will require substantial computational effort. The objective is to keep future simulations in acceptable ranges of computational costs, depending to future use of the model (parameter exploration, protocol optimization, etc.).

#### Step 2: cell types and attributes

Once the environment and its governing rules are set, we define the cell types that will intervene in the modeled processes. By cell type, we refer to cells that will have different parameters, behaviors and interactions in the simulation. We expect them to mostly correspond to known biological classifications. The objective is to segment the different behaviors and clarify their description in the following steps. Additionally, the attributes (i.e. internal characteristics) of the cells are described. Attributes can be cell radius, division counter, various protein production/consumption capacities or any other value necessary for the development of the model.

#### Step 3: global cell behaviors

Once the cell types defined, we then describe the behaviors of the cells. To this end, we use state-transition diagrams (Fig 2.A). These diagrams can be easily translated into code and have the advantage of being easily understandable for non-specialists. They are widely used in software engineering to understand customer needs in the Unified modeling Language (UML) methodology [20, 21]. In this diagram, each rounded rectangle describes a state a cell can be in and each arrow a transition from a state to another. This aims to abstract the different phenotypical expressions of the cells. The initial point (fulfilled black circle) describes the initial state a cell is in at its birth or its arrival in the simulation frame. At each step of the simulation, a cell behaves according to its state (see Step 4), its inner properties can change depending on their actions and environment. At the end of a simulation step, the cell changes of state if the conditions (labeled on the arrow) are respected. The final point of the diagram (fulfilled surrounded black point) corresponds to its removal from the simulation.

**Fig 2.**
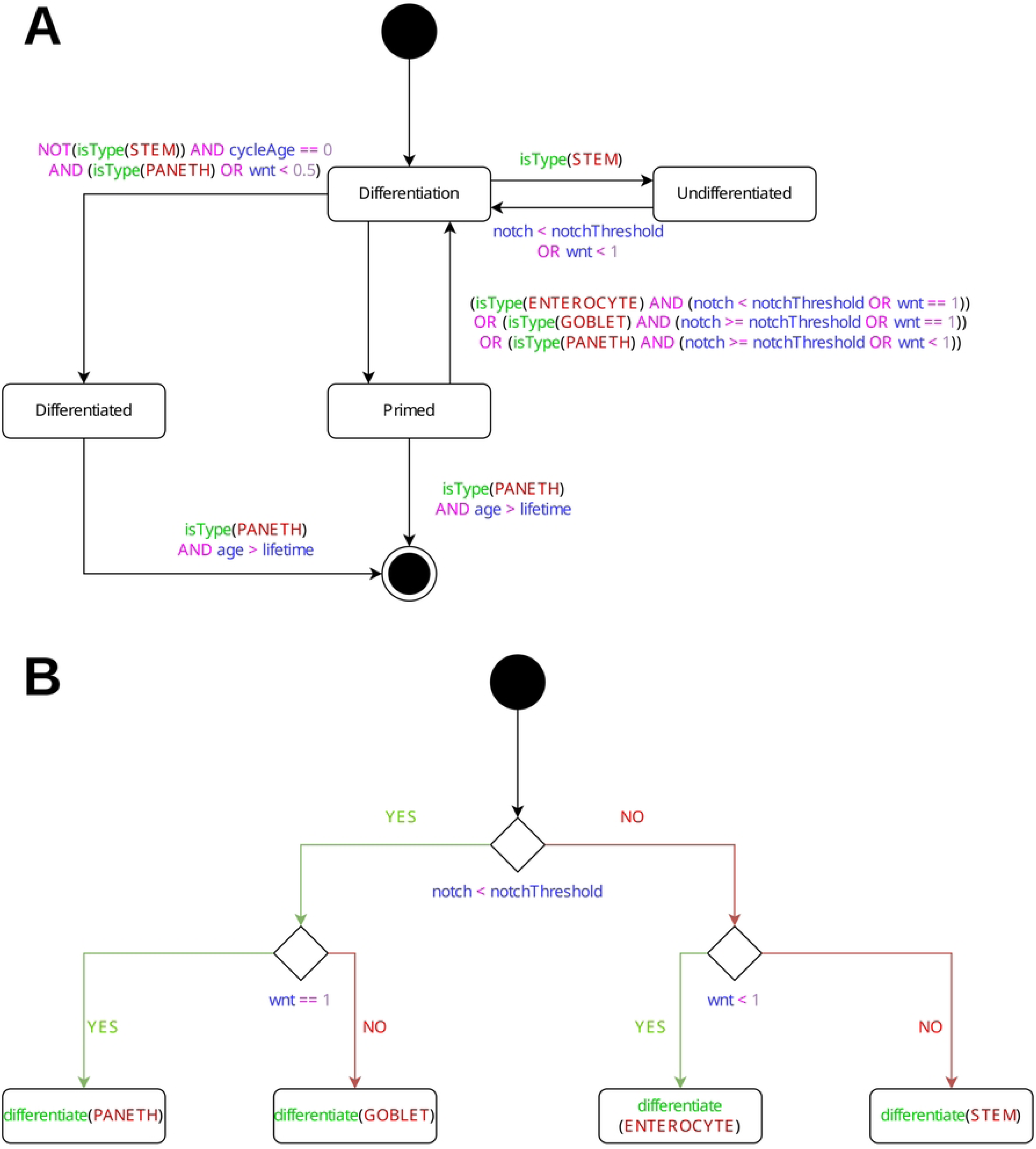
State-transition and activity diagrams. State-transition diagram (**A**) and activity diagram (**B**) extracted from the third study case model of the results section (see this section for more details). The activity diagram corresponds to the differentiation state of the state-transition diagram. **A** : Depending on the molecule quantities the cell is subjected to, a stem cell can differentiate in 3 different types (enterocytes, paneth or goblet) or stay undifferentiated if the conditions are not met. A primed cell can still differentiate if their condition changes. After meeting the conditions to enter the differentiated state, the cell can not enter the differentiation state anymore. Primed and differentiated cells can die if they exceed their lifetime without dividing or going out the crypt. **B** : Cells in the differentiation state can differentiate in 3 different type or stay/return stem cells. If the notch quantity is lower than the notch threshold, the cell can become one of the secretory types depending on the wnt quantity (paneth or goblet) otherwise if the wnt quantity is lower than 1 it becomes enterocytes instead of a stem cell.

#### Step 4: cell actions

We then determine the action sequence the cell follows in each state. For this purpose, we use activity diagrams [22, 23](Fig 2.B). In this diagram, rectangles describe basic actions (such as growing, randomly moving or dividing) and diamonds binary conditions. The arrows, for their part, describes the sequence of actions each cell will follow during one time step. The main objective of this diagram is to visually program the behavior of the cell. Modelers must though implement the inner functioning of each basic action (i.e. the rectangles).

#### Step 5: experimental protocol

Finally, the experimental protocol must be designed. Here we describe the initial conditions and the external events that will be triggered at different times of the virtual experiments. This draws the parallel with real wet lab protocols usually designed by the biologists prior to any experiment.

As in step 4, activity diagrams are used to visually describe the initialization of the simulation (physics, cell seeding, molecular gradients, etc.) and in-simulation events (cell addition, drug injection, etc.).

### ISiCell: a web platform to interactively build and explore *in silico* models

While we argue that the methodology presented in the previous section allows a better involvement of the biologists in the design process and therefore a better understanding of the produced models, it is necessary to build programs as fast as possible in order to catalyze this interaction. To this end, we propose a web-based modeling platform which allows (1) to draw diagrams from the proposed methodology, (2) to automatically build executable simulation from the methodology schema and (3) to explore and analyze the created model. To this end, the platform disposes of three main tools:

- **ISiCell Builder**, a schema-to-code tool which generates C++ code from drawn diagrams and compile it to build an executable simulation,
- **ISiCell Viewer**, a 2D/3D visualization tool in which parameters and experimental protocols can be set up, the corresponding simulation can be explored both via a graphical representation of the cells and environment and via customizable plots and statistics,
- **ISiCell Explorer**, an innovative parameter exploration tool that allows to visually explore various sets of parameters and their impacts on the simulation kinetics.

This platform is accessible and testable at this url: https://isicell.irit.fr.

### ISiCell Builder: a schema-to-code tool to support the methodology

The main objective of ISiCell Builder is to interactively set up the models by selecting models of environment and by drawing the state-transition and activity diagrams from the methodology. Fig 3 shows the graphical interface of the builder.

**Fig 3.**
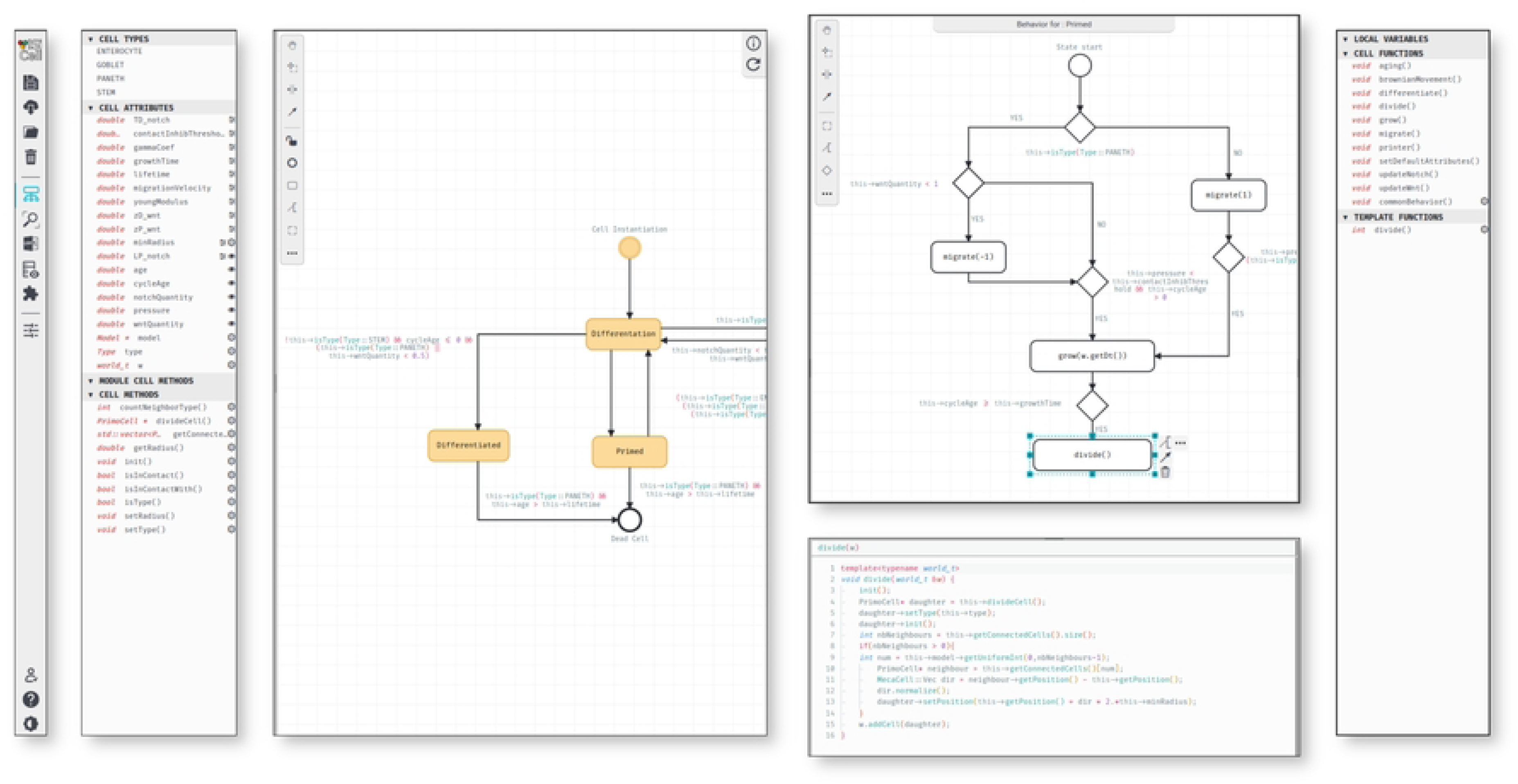
ISiCell Builder interface. Cell Builder is the main tool for prototyping the models alongside biologists with an easy-to-follow interface. This figure represents the builder view for the before mentioned murine intestinal crypt model. From left to right, the main menu bar allowing to navigate through the different tools; the attributes, methods and types lists where you can add new type to the simulation and new characteristics to the cells; the state diagram, specifying all the states any cell can go through and the transitions between these states; the primed state activity diagram with the divide C++ written function code below; the created functions and generic templates which can be dragged and dropped in the activity diagram.

To set up the environment, the platform already implements various physical (Mass-spring-damper, Hertzian, variable density grid, etc.) and diffusion models that can be selected and added in one-click. Thanks to a plugin-oriented architecture, new modules can be added to the platform for specific needs to simulate new type of environment, specific behaviors or high-level statistics.

The main panel of ISiCell Builder allows to build the behavior of the cell by drawing the state-transitions diagram of the global cell behavior and the activity diagram of the specific behavior of each state. To implement these specific behavior, a C++ editor is also available for the modeler. Pre-implemented functions such as divide, migrate, grow are also available and can be added as is or modified in the activity diagram.

Once designed, the model can be automatically translated in code in one click. The simulated environment is set up by plugging in the selected modules to the program. The state-transition diagram is then transformed into two functions:

- The behavior function executes the code associated to each state. The code of each state is generated based on the flow described in the activity diagrams: each box of these diagrams corresponds to a function call which code has been implemented in the C++ editor during the model design.
- The transition function updates the state of the cells depending on their current state and the condition implemented on the transition arrows.

All in all, the generated code is guaranteed to be exactly equivalent to the diagrams drawn and is then compiled to obtained an executable program.

### Interactive visualization and exploration of the generated model

ISiCell Builder is the main tool of the platform enabling fast prototyping following our methodology. In order to quickly have qualitative feedbacks from the biologists and to improve the biological coherence of models, the platform disposes of visualization and exploration tools. These tools have been designed with the objective to involve biologists in the exploration of the model and collaboratively explore sets of parameters to validate the coherence of the system, test simple hypotheses or perform a summary calibration of the model.

#### ISiCell Viewer

Agent-Based models easily allow for visualization. Visualizing a prototype helps the biologist to qualitatively validate the coherence of the model. Thanks to their knowledge of the biological reality, biologists can provide a critical insight on the individual and global behavior of the cells. Thus they can help correcting and/or rapidly improving the model.

ISiCell Viewer (Fig 4) has been developed for visualizing and summary analyzing the code generated by ISiCell Builder. It offers the possibility to launch simulations with different parameters and observe directly their impact on the model or test simple hypotheses. The idea is to use visualization to refine the model in ISiCell Builder and going back to ISiCell Viewer to visualize the impacts of the modifications until the biologists find it adequate.

**Fig 4.**
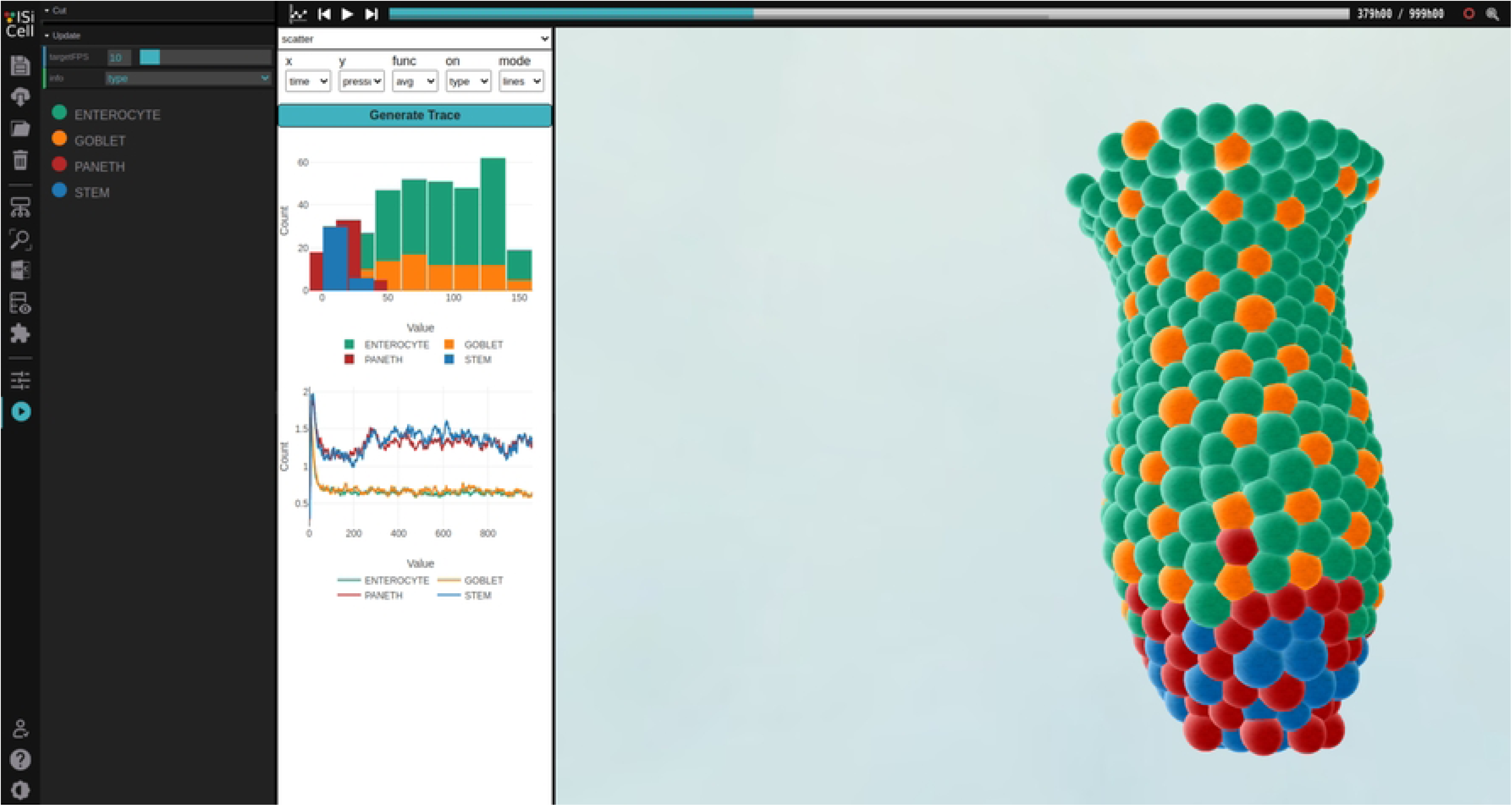
CellViewer overview on a intestinal crypt model. The upper bar enables to go back and forth in the simulation. On the left side panel coloration can be changed. The white panel allows for dynamic generation of plots.

ISiCell Viewer possesses several tools to help the qualitative evaluation of a running simulation. It provides a graph panel allowing to plot most of the simulations’ attributes about the population, spatially or temporally. It also allows to move to any time in the simulation, visualize different colors on cells depending on their attributes, create cross sections, etc.

Once qualitatively approved by the biologists, the model’s parameters needs to be explored to be calibrated on experimental data and/or evaluate the impacts of parameter modifications on the simulated kinetics. Most of the time, the exploration is done by the modeler with no link with the domain expert, potentially leading to non-realistic sets of parameter. Together with the visualization tool, we propose in the ISiCell platform a parameter exploration tool to interactively sample parameters and dynamically visualize their impact on simulated kinetics.

#### ISiCell Explorer

ISiCell Explorer enables to launch several simulations with different parameters in parallel and draw personalized plots for each of them. The goal is to offer a primary sensitivity analysis tool to have a qualitative insight on the impact of chosen parameters on the model.

This tool is based on editable Python scripts to generate plot and launch simulations. So that the script has access to the data of the simulation at each time step, the code is specifically wrapped and recompiled. This wrapping allows to directly and simply interact with simulations with a programming language, Python, adequate for data analysis. As it is written in Python, the tool also offers to easily load biological data in order to plot them with the simulation for comparison.

ISiCell Explorer allows for exploring one or more parameters at a time between selected range. The simulation will be launched with variation on the selected parameters following a Latin Hypercube Sampling [24, 25]. The generated results can be selected to relaunch the exploration using another set of parameters as a basis. Thus it is possible to reiterate until finding an interesting set of parameters. The interface offers to construct a tree of each iteration allowing for going back and forth trials (as shown in Fig 5).

**Fig 5.**
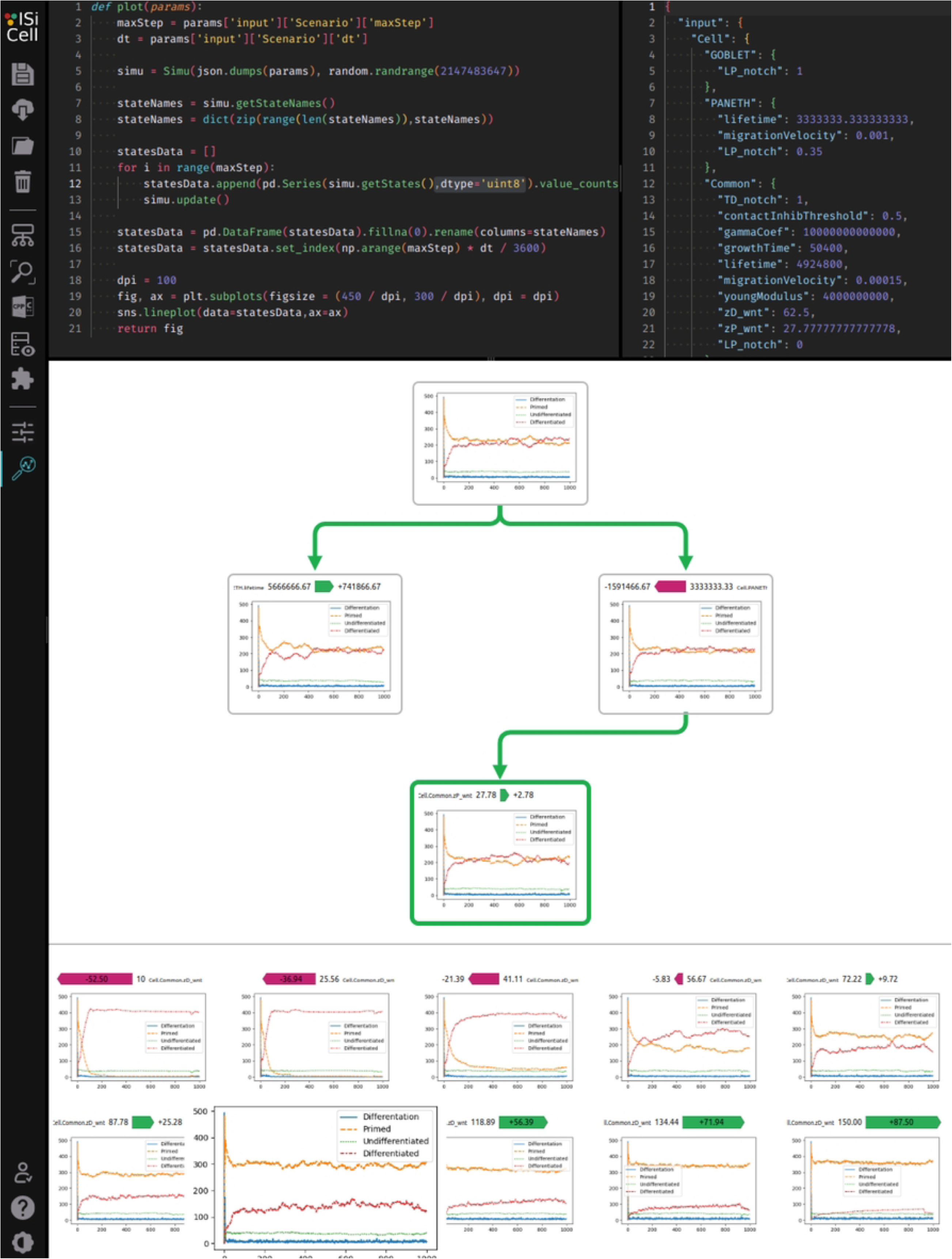
CellExplorer view after a few parameter explorations. The upper left panel contains the python script launching the simulations and generating the plots. The upper right panel contains the inputs values in json format.

This tool enables to easily explore the input space and observe their impact on the population dynamics. This tool can be very powerful to test simple hypotheses or to verify that the model follows already known biologic dynamics. For the biologists, it could lead to a better understanding of the inner mechanisms of the cells and their impact at the population scale.

## Results

In order to illustrate the range of possibilities of the platform, various models both based on the literature and from workshops we organized will be presented in this section. Table 1 summarizes them all :

**Table 1.** Model case studies summary

| Case study | Type of environment | Used modules | Short description |
| --- | --- | --- | --- |
| Cancerous spheroid | 3D continuous | Mass-Spring physics, Oxygen Diffusion, Spheroid Manager | Growth of cancerous spheroid in a 3D environment involving oxygen diffusion and cell cycle dynamics |
| CTL interaction with tumors | 3D continuous | Hertzian contact physics, 3D environment | Monolayer tumor growth control with sequential addition of Cytotoxic T-Lymphocytes |
| Intestinal crypt | 3D continuous | Hertzian contact physics | Renew process of a murine intestinal crypt |
| Wound inflammation | 2D discrete | 2D Diffusion, 2D Grid | Inflammatory response to a wound in a toric discrete 2D muscular tissue |

Summary of all the model case studies presented in this section.

Our methodology offers the advantages of making models easier to understand and to reproduce even for people who didn’t work on them thanks to its diagrams.

### Case study 1: cancerous spheroid growth and cycle dynamics

Spheroids have been used by biologists to study cancerous tumor for decades [26]. They have the advantages of reproducing the first stages of a tumor formation before vascularization. Moreover, they have been widely used for antitumoral treatment preclinical evaluation. Mathematicians and computer scientists provided another level of abstraction by in silico modeling spheroids [27–29]. For the last decade, modeling 3D cell culture has become quite common.

The objective of this in silico model was to get a realistic behavior reproducing the proliferation gradient inside spheroids and associated cell cycle dynamics as these parameters impact on the response to antitumor molecules. The development performed during workshops allows to define the state-transition diagram and the rules for corresponding behavior together with biologists taking into account previous collaborative works on cancerous cells, study of response to microtubules targeting agents inoculation and impact on cycle dynamics. [30–33]. The interest of the approach used here was to obtain very quickly a first model with an instantaneous visualization using the viewer allowing to directly make correction for different parameters as seen in Fig 6. The other point is linked to the graphical representation of the rules of the model allowing biologists to much better interact with the model than regular code. This is an essential aspect of acceptation and implication of the interdisciplinary work by biologists.

**Fig 6.**
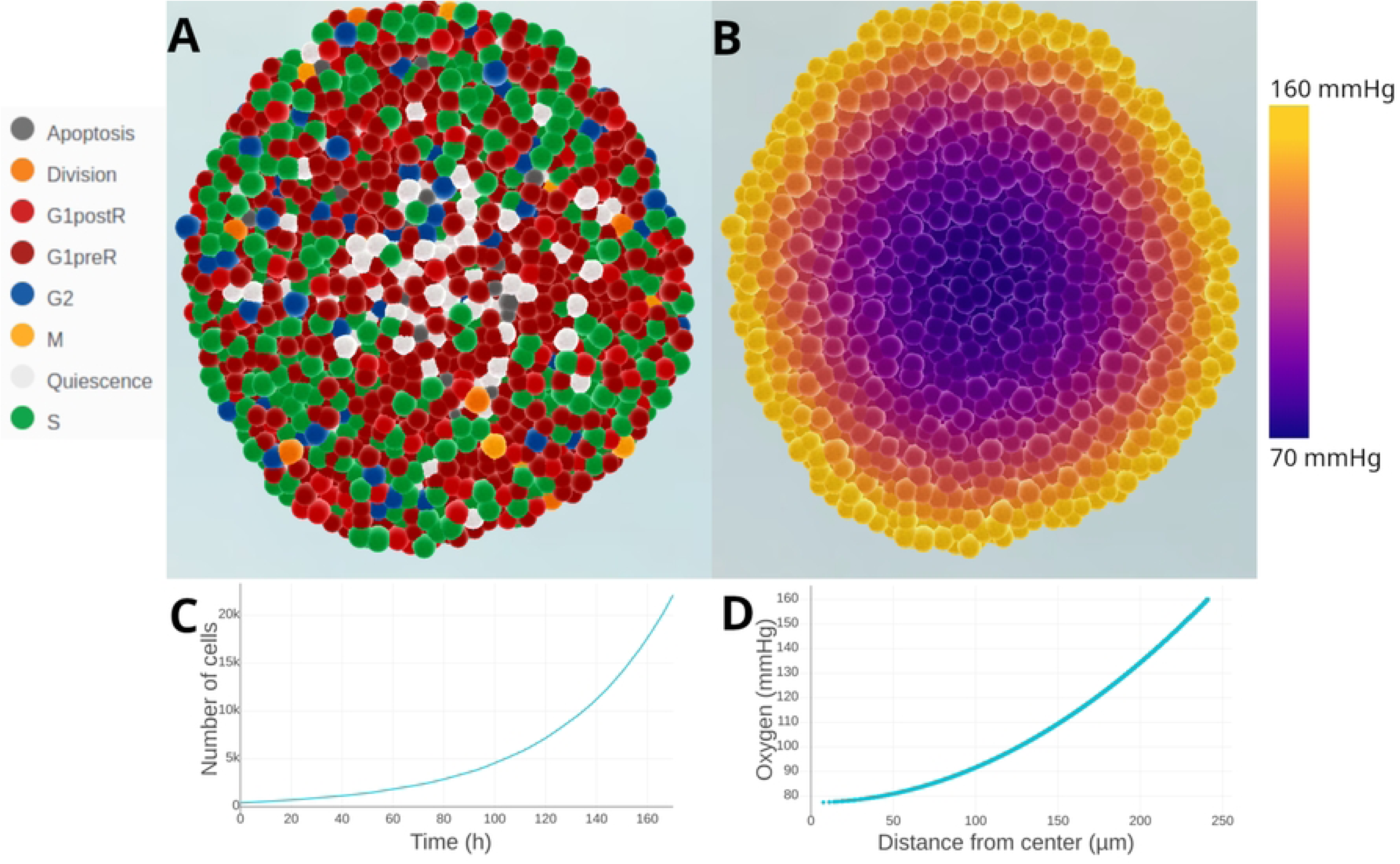
Cut visualizations and ISiCell Viewer plots of an above 20 000 cells spheroid. (A) displays the state of each cell, cells are less proliferative at the center. (B) shows the oxygen gradient in the almost 250 µm spheroid. (C) describes the exponential growth of the spheroid in terms of number of cells. (D) illustrates the oxygenation depending on the distance to the center. Without experimental data, biologists can still qualitatively validate the coherence of the simulated spheroid using the ISiCell viewer tool. This model is available here: https://isicell.irit.fr/app.html?demo=Spheroid.

This model involves a mass spring physics module [34] to obtain the shape of the spheroid. Each cell is linked to its neighbors in the same way as if there was a spring connecting their two centers. An oxygen gradient [35] module is also used to manage the oxygen quantities in the spheroid. The behavior is split between the different phases (as shown in Fig 7) of the cell cycle whose durations are settable to match specific cycle depending on the studied lineage. It offers the possibility to test different population dynamics by playing with the cycle rules and parameters. For example, the influence of the elongation of the G1 phase depending on the oxygenation or the impact of the quiescence rate can be tested. In the same way, inoculation of molecules that disturb the cell cycle can be tested to reproduce their effects on the culture.

**Fig 7.**
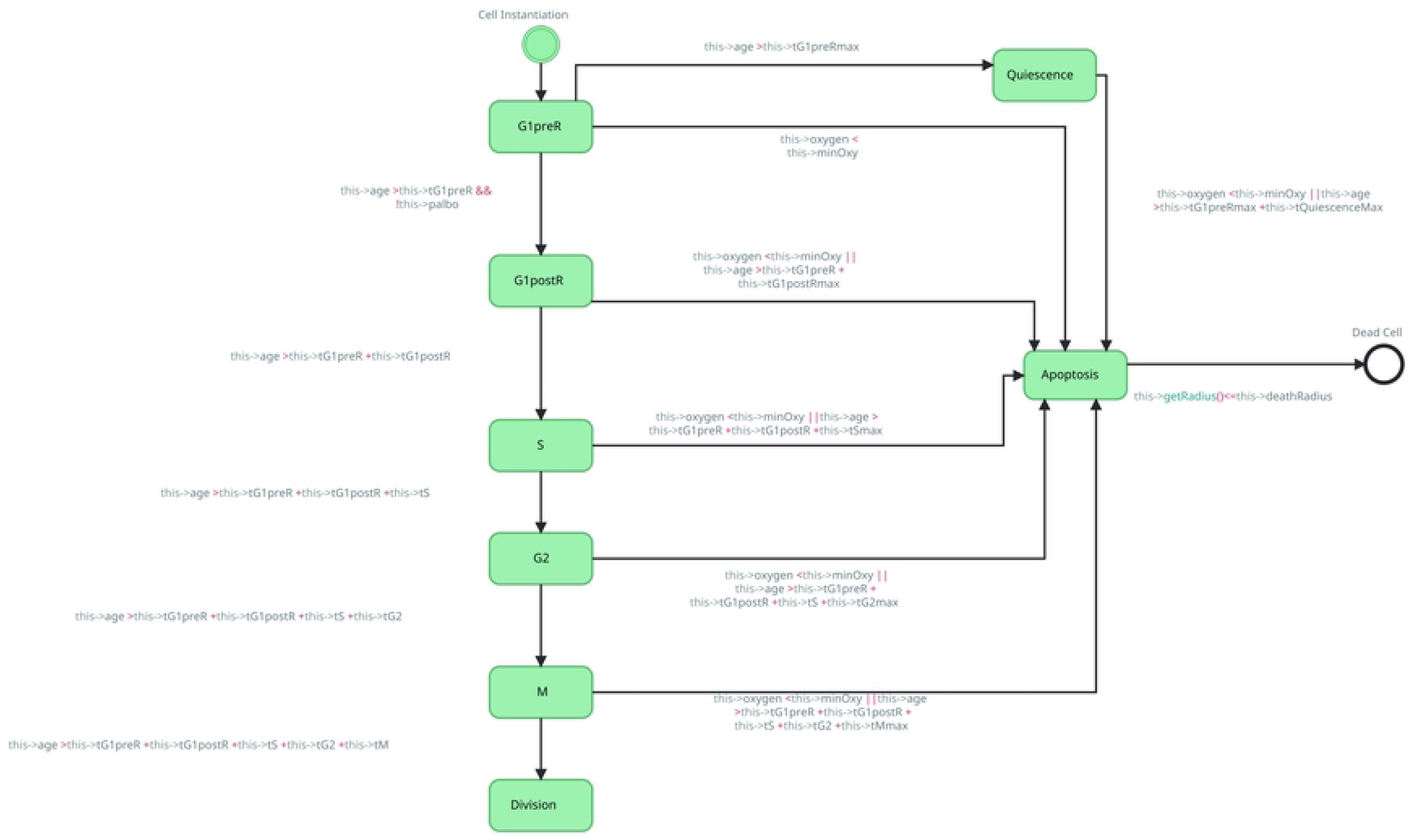
Spheroid State Transition Diagram. The behavior is split between the different state of the cell cycle. Cells can also enter a quiescent state or apoptosis. Cells are removed from the simulation after shrinking to a minimal radius.

This model is also interesting to benchmark the computing time of the platform depending on the number of cells. The number of cells grows exponentially and likewise the computing time needed to update the behavior of each cell and especially to calculate their new positions with the physics module. In the Fig 8, a computing time comparison is made between the three mode of use of the generated simulation showing that the visualization and the wrapped Python code are respectively averagely 2.1 and 1.3 times slower than the C++ standalone.

**Fig 8.**
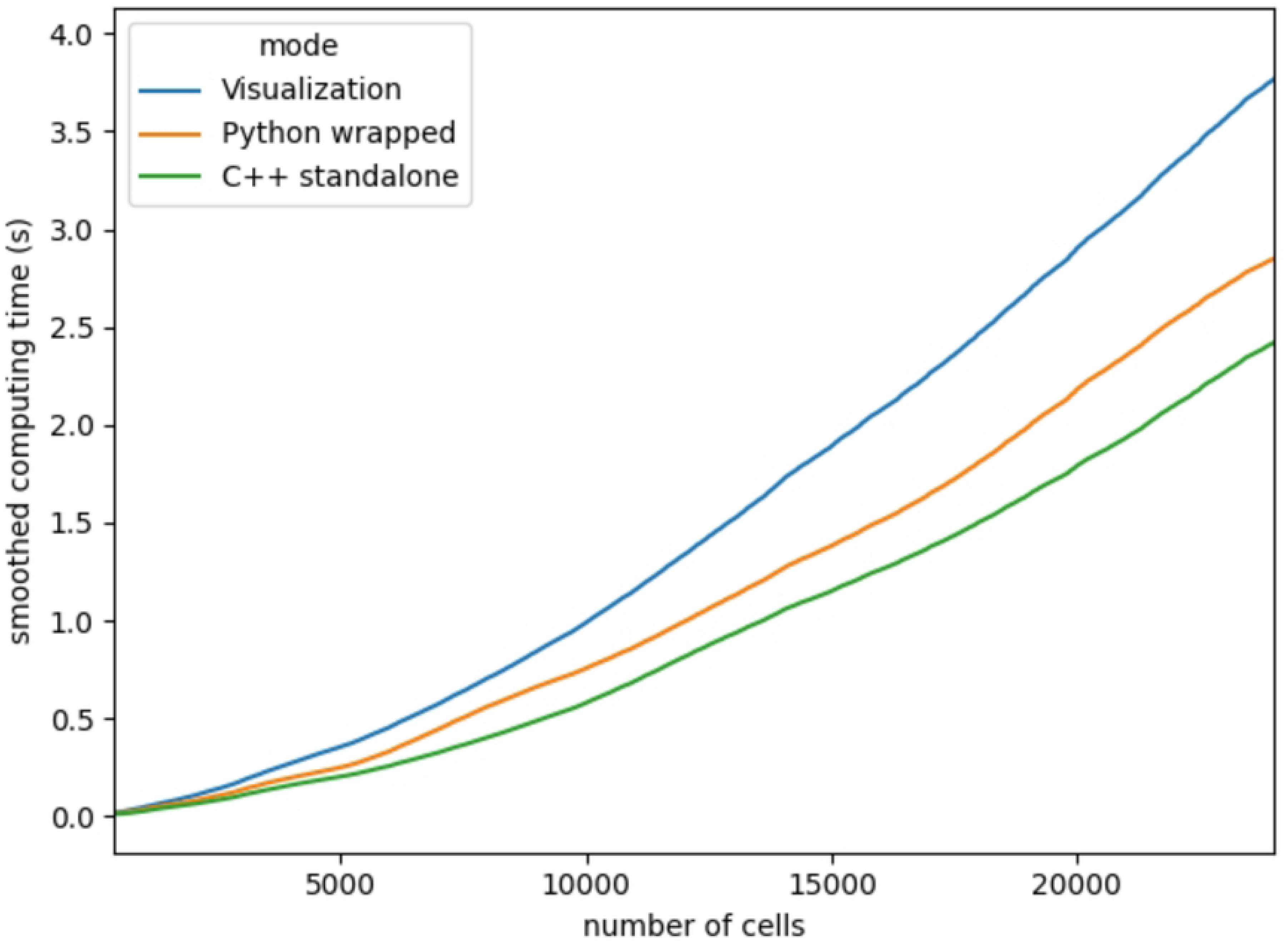
Compared computing times. Smoothed computing time plot showing the time needed to compute a step of simulation depending on the number of cells. The computing time was calculating with the same seed on a 200 hours spheroid simulation with a 2.70GHz Intel® Core™ i7-10850H CPU and 32Go of RAM. Each simulation step represents 10 minutes in the simulation. The visualization and python wrapped execution respectively multiply the computing time by an average of 2.1 and 1.3

### Case study 2: sequential adjustment of Cytotoxic T-Lymphocytes to control tumor growth

In silico models are more and more often used for therapeutic protocol exploration [36–38]. As a matter of fact, a well calibrated model constitutes a predictive tool that can help exploring the different protocol possibilities quicker and more efficiently. Thus they can provide optimized protocols answering specific constraints.

This second case study was developed during workshops and is based on previous works on immune treatment of cancer cells culture [39]. It focuses on the 2D growth dynamic of cancer cells in presence of Cytotoxic T-Lymphocytes (CTL) cells which are sequentially added.

Cancer cells grow until the end of their cycle before dividing. If they get in contact with an active CTL, they enter a defensive state. In this state, they have a probability to inhibit CTLs in their surroundings at each step of simulation. If there are no more CTLs attacking, they go back to their cycling state or else they have been killed by CTLs and are removed from the simulation. Active CTLs randomly roam in the simulation environment until getting in contact with a cancer cell and enter an offensive state. In this state, they attack the cancer cells with which they are in contact until getting inhibited or killing them. After killing cancer cells, active CTLs go back to their roaming state. If they got inhibited by a cancer cell, CTLs enter an inhibited state where they indefinitely roam without attacking anymore. The corresponding state transition diagram corresponds to the Figure 9. The hertzian physics module [34, 40] was used to shape this 2D continuous model and manage the cells movements. You can see an example protocol of the model in Figure 10.

**Fig 9.**
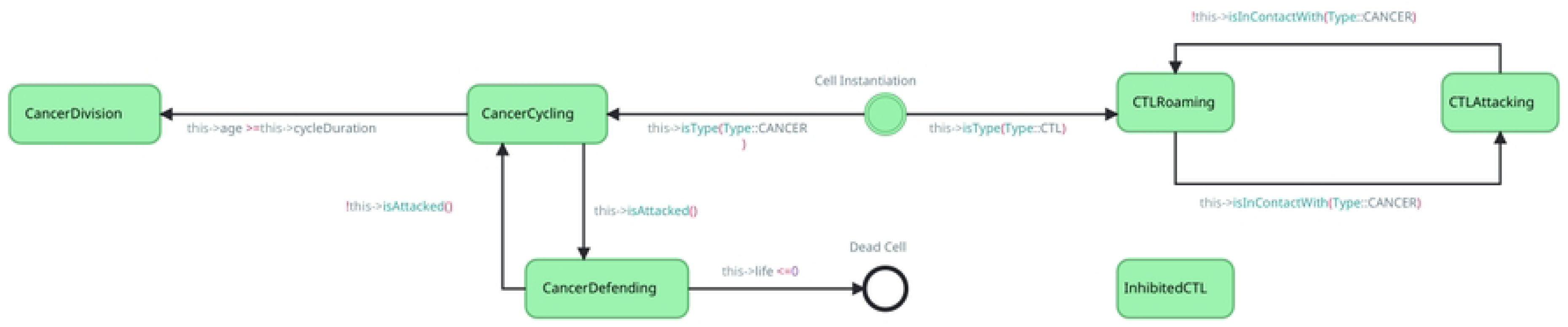
State transition diagram of case study 2. The behavior is split between the CTL and the cancerous cells.

**Fig 10.**
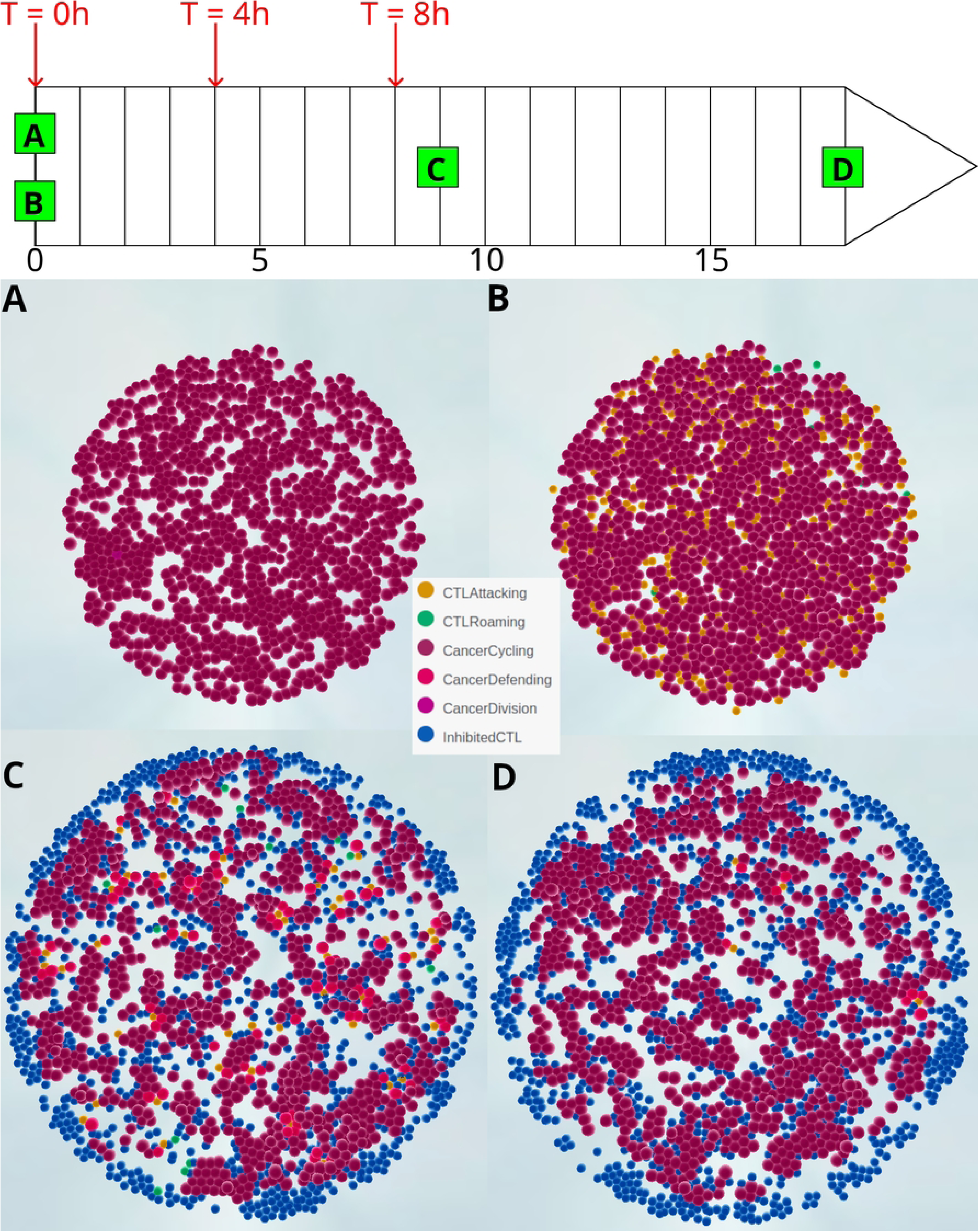
Series of screenshots of a 18h simulation with 3 CTL injections and 1000 initial cancerous cells. The protocol is displayed above with 3 injections at T=0h, T=4h and T=8h for a total of 1000 CTLs. (A) shows the initial cancer cells disposition followed by the first CTL injection (B). (C) shows the simulation after 9 hours and three CTL injections and (D) represents the state of the simulation at the end. This model is available here: https://isicell.irit.fr/app.html?demo=Immunology

The protocol design tool of the platform offer the possibility to set several CTL injections during the simulation in order to reproduce a clinical treatment on the cell culture. The newly built model succeeded in reproducing the population dynamics of the original study as seen in figure 11. These in silico experiments suggest that dividing the CTL injections is more efficient for controlling the tumor growth.

**Fig 11.**
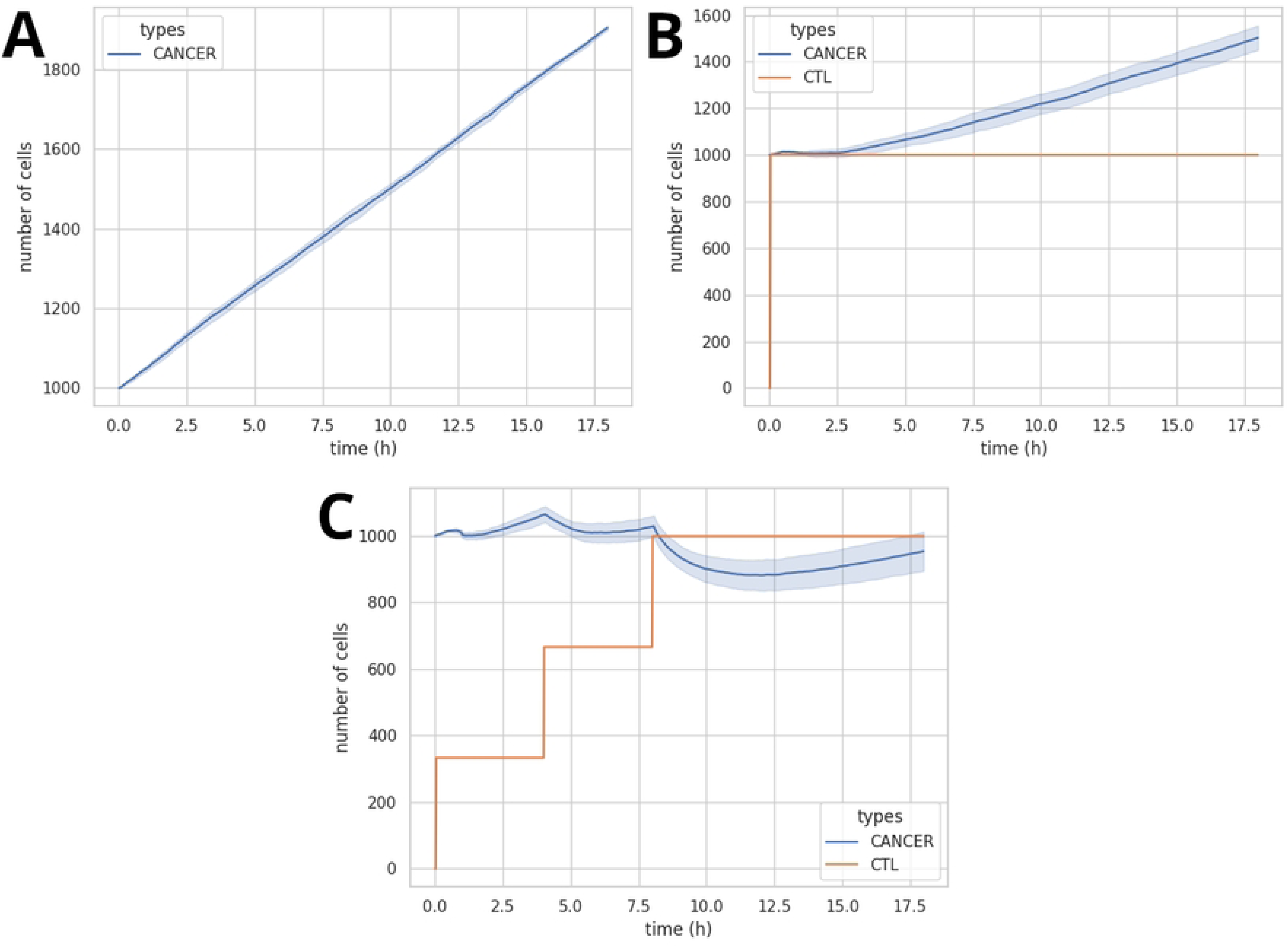
Population dynamics graph generated depending on the CTL injection protocol. (A) control population without CTL (B) 1:1 injection protocol with one injection at T=0h (C) 1:1 injection protocol with 3 equal injections Results show that dividing the injections improve the control of the tumoral growth

### Case study 3: dynamic model of stem cell and tissue organization in murine intestinal crypts

The intestinal crypt model has long-standing history in computational modeling [41]. It has drawn the interest of modelers for decades because of the complexity of the mechanisms involved in the renewal of cells and their spatio-temporal differentiation in the structure.

The third application presented in this paper is based on a model published in 2011 by Buske et al. [42]. In this model, 4 cellular types (stems, paneths, enterocytes and goblets) are interacting to continuously maintain the intestinal crypt. This renew process is driven by two main conditions: the Wingless and Integration site (Wnt) gradient (higher at the bottom and lower at the top) and notch quantities in the crypt. These two molecules regulate the differentiation of the cells. For simplification, the wnt quantity has a value of 1 under a *z_P_* position, a value of 0 above a *z_D_* position and a value of 0.5 inbetween. The *z* = 0 position corresponds to the bottom of the crypt. In the same way, instead of a mesh with which the cells interact, axis forces are used to maintain the crypt’s shape.

Stem cells, mostly found in the bottom of the crypt, can divide when physical pressure is lower than a given threshold. To divide, they grow in size for a given period of time before actually dividing. Stem cells can also go in a differentiation process if their notch quantity is not high enough or if they enter a upper part of the crypt where wnt is lower than 1. If the notch quantity is not high enough, they can differentiate into the paneth type if they still are at the bottom of the crypt where the wnt quantity is higher or become goblets if they are above the *z_D_* position. These two types are called secretory because they increase the notch quantity in the cells with which they are in contact. If the notch quantity is high enough stem cells can differentiate into enterocytes when quitting the lower part of the crypt.

Freshly differentiated cells can enter a prime state where they can divide or differentiate once again if their notch or wnt quantities do not match their type. If goblets or enterocytes reach the upper part of the crypt, where the wnt quantity is the lowest, they cease to divide after a last division and enter a differentiated state in which they will not be able to replicate or differentiate anymore. Paneth cells tend to immediately enter the differentiated state if they are at the bottom of the crypt.

This complex process is visually represented by the state-transition diagram presented in Fig 2.A. This first study uses the preimplemented hertzian contact physics [34, 40] module which allows to maintain the crypt shape and to facilitate cell migration.

All in all, our methodology associated to the ISiCell platform allowed to simply reproduce Buske et al.’s model. We showed the capacity of the platform to represent very well described model and to reproduce kinetics comparable to the one obtained in the original authors work. The main contribution of our platform in this case study lies on graphical programming of the model as presented in Fig 2.A and the direct graphical visualization of the on-the-fly generated simulation, as presented in Fig 12. As depeticted in this figure, the cell behavior can be easily read by user outside of the modeling domain. Additionally, such a diagram can be added as is in a publication to improve the replicability of the work: such a state-transition diagram can be simply recoded to obtain an equivalent model.

**Fig 12.**
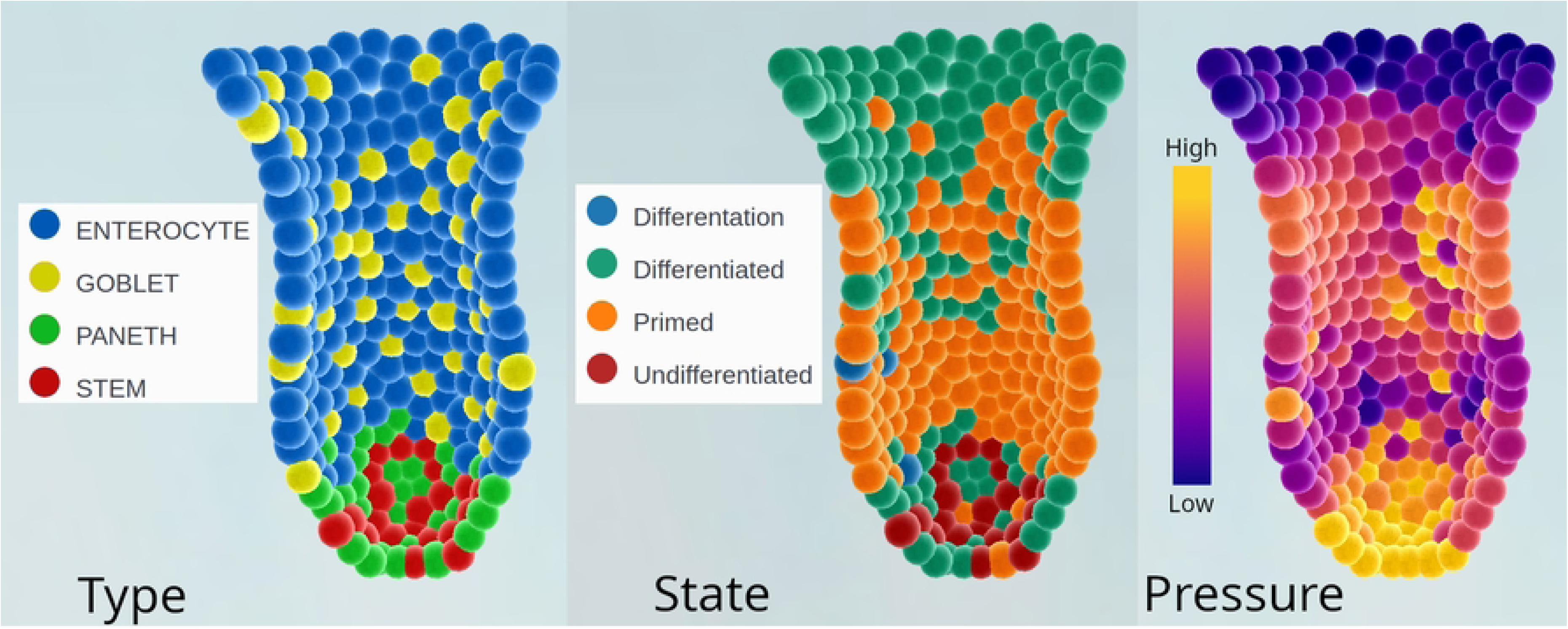
Intestine Crypt cut visualizations. From left to right, the type distribution in the crypt, followed by the state distribution and the local pressure in each cell. The visualization helped to qualitatively validate the model by comparing with the expected results. This model is available here: https://isicell.irit.fr/app.html?demo=Intestine_crypt.

### Case study 4: muscle wound inflammation

The immune response is a complex phenomenon involving different cell types and chemical transmitters dynamically affecting each other. This intricate dynamic has been the subject of several computational models [43–45] for many years.

The fourth case study is based on [46] which describes a 2D discrete toric grid of individual muscular cells (IMC). While bacteria grow on IMCs, they cause an inflammatory response through chemical messages and chemotaxis. The original model was developed using NetLogo [47].

IMCs are initially in a resting state. They enter the activated state if the amount of bacteria, reactive oxygen species (ROS) or damage associated molecular pattern molecules (DAMP) are higher than their respective threshold and start producing interleukin-8 (IL-8). They can go back to their previous state if the 3 previously mentioned quantities decrease down under their threshold, the IMC will then stop emitting IL-8. However if one of this 3 quantities increase above their higher second threshold, they become damaged IMCs and start to be attacked (represented by the loss of life points depending on the quantities of ROS, DAMP and bacteria) until the cell enters apoptosis (when they have no life points anymore) or is phagocytized by a macrophage. When entering the damaged state, an IMC emits a large amount of DAMP. In the damaged and apoptosis states, damaged IMCs produces a little amount of DAMP. This DAMP secretion was not specified in the original paper but seemed necessary to guide macrophages in removing damaged and dead cells. A high quantity of ROS can also remove a damaged IMC in apoptosis state from the simulation.

Neutrophils enter the simulation through blood vessels and get activated by the IL-8 quantity in their position. In their initial state, they follow the IL-8 gradient and produces monocyte chemoattractant protein 1 (MCP-1), interferon gamma (IFN-*γ*) and ROS until being in the same position as a damaged IMC, being confronted to not enough IL-8 or getting too old (meaning they are moving in the grid without finding a damaged IMC for too long). In the two latter case, the neutrophil is removed from the simulation. In the first case, a neutrophil phagocityzes the bacteria in its position unless another neutrophil or a macrophage did it just before. In the second case, the neutrophil will go back to its previous state. Otherwise it removes all the bacteria in its position and emits a great amount of ROS before becoming pus. In their state, pus only emit a little amount of DAMP for the same reason as the damaged IMCs.

Undifferentiated macrophages enter the simulation through blood vessels in the same way as neutrophils. In this state, they follow the MCP-1 gradient until becoming differentiated macrophages or getting too old (meaning they are moving in the grid without differentiating for too long) and being removed from the simulation. The differentiation is triggered by a high enough amount of IFN-*γ* or DAMP. Differentiated macrophages start emitting tumor necrosis factor alpha (TNF-*α*), following and absorbing DAMP. If there are bacteria, damaged IMC or pus in its position, a macrophage phagocytizes them by removing them from the simulation. If the amount of DAMP is low enough and the macrophage is near a healthy IMC without damaged IMC or pus nearby, it enters a resolutive state where it starts producing Interleukin-6 and -10 (IL-6 and IL-10) and stops emitting TNF-*α*, moving and absorbing DAMP. They can go back to their previous state if the conditions needed to enter the resolutive state are not respected anymore. This state was not mentioned in the original article but seemed necessary to reproduce the healing dynamics.

Myoblasts also enter the simulation through blood vessels. They start by following the IL-6 gradient and get activated in the presence of this same molecule. When activated, myoblasts secrete IL-6 in presence of a sufficient quantity of IL-10. They continue to follow the IL-6 gradient until they found a position without bacteria and low enough DAMP amount adjacent to a healthy IMC or another myoblast. In this condition they start their fusion to restore the damaged tissue. In this state, myoblasts do not secrete any molecule anymore. After a certain time, myoblasts enter a differentiation state and become new healthy IMC.

The last type of agent in this model is the blood vessel type. They are homogeneously disposed in the grid at the initialization. They have only two states : open or thrombose. Vessels stay open as long as there are an healthy IMC in their position and at least 2 other healthy IMC in their direct neighborhood. In this state, they can bring neutrophil, macrophage or myoblast in the simulation depending on the quantity of dedicated molecules. Neutrophils are called by a sufficient amount of IL-8 or TNF-*α*, undifferentiated macrophages by MCP-1 or DAMP and myoblasts by IL-6. After getting at least one of this previously mentioned cell, a blood vessel has to cool down for a certain time. This cool down mechanism was not mentioned in the original article but seemed necessary to regulate the number of cells entering the simulation.

In this case study, bacteria are not treated as agents but as local values in the grid with their own management. There is two types of bacteria, the regular ones that can grow on damaged IMCs and the virulent ones that multiply slower but can grow on healthy IMCs. The first type produces a virulent bacteria for one thousand regular bacteria. When attaining the carrying capacity for one position in the grid, bacteria start multiplying on the direct neighboring positions if there is a damaged or healthy (only for the virulent type) IMC. Their number also decrease depending on the ROS quantity; virulent bacteria also have a better resistance to ROS than regular ones.

In order to reproduce the diffusion of molecules in the grid, the platform proposes a module to manage 2D diffusion which is here coupled with a 2D discrete environment for the cells. In order to confirm the coherence of the cells behaviors, we use ISiCell Viewer which enables to qualitatively reproduce the dynamics of the original article (as shown in Fig 13). To calibrate the model we used the ISiCell Explorer tool to qualitatively match the population dynamics graphs of the original paper (as shown in Fig 14).

**Fig 13.**
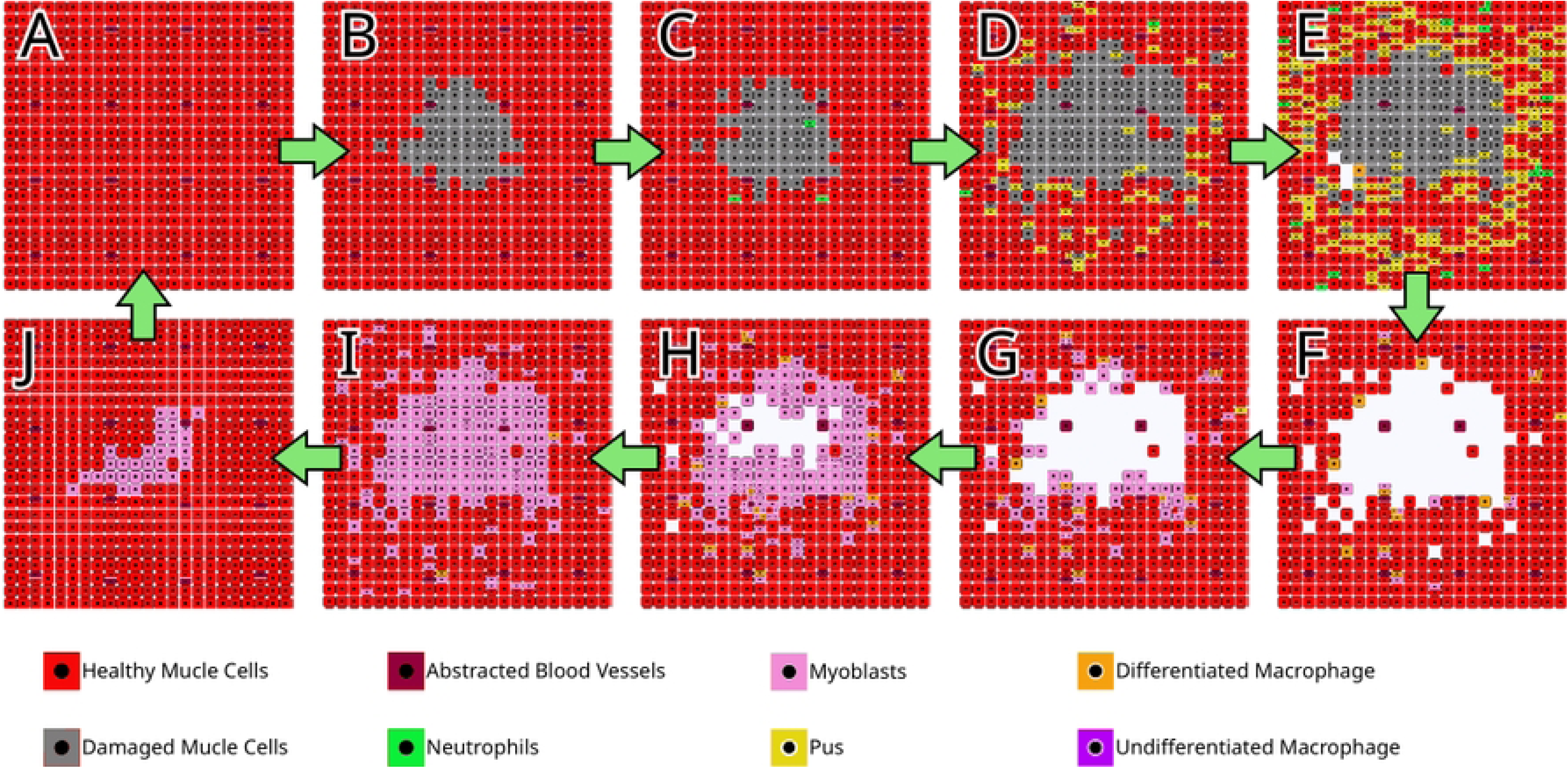
Series of screenshots of a single representative run demonstrating successful healing reproducing results found in the original article. (A–J) show the progression of the tissue from healthy to restored after damaged. CellViewer helped to obtain this qualitative reproduction. This model is available here: https://isicell.irit.fr/app.html?demo=Muscular_inflammation.

**Fig 14.**
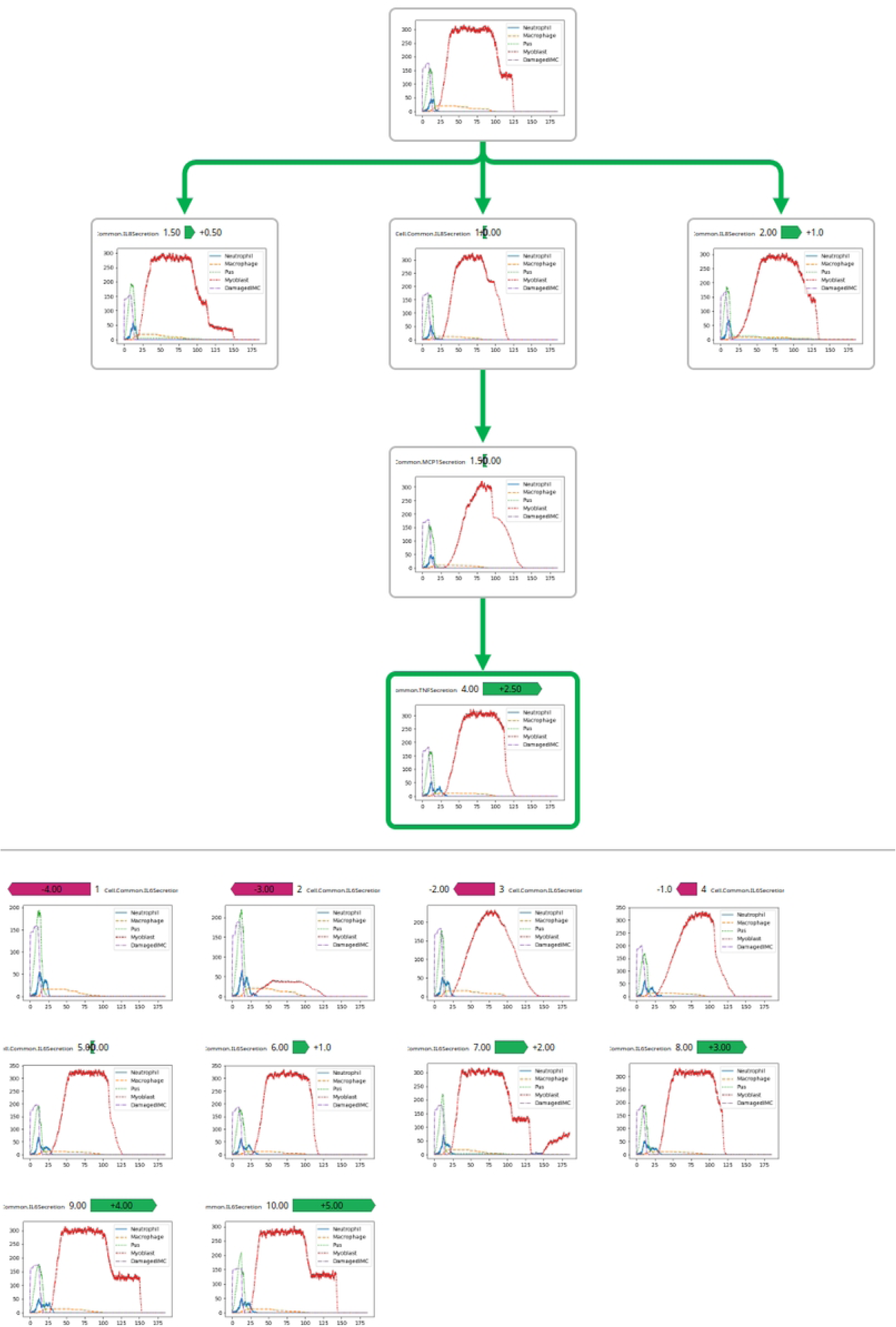
Parameter exploration of the muscle wound model using ISiCell Explorer. This tool revealed to be useful to study the impact of each molecule secretions on the type dynamics helping us to reproduce dynamics from the original article.

On the basis of Gopalakrishnan et al.’s wound healing model, we obtained a similar inflammation model reproducing complex dynamics comparable to the one provided in the original paper. The complexity of this models is reflected in its state-transition diagram (see Fig 15) resulting from the application of our methodology. However, this diagram allows for a better understanding of the different type behaviors. Moreover the explorer and viewer tools make this model an interesting toy model to study complex interactions and directly observe the impact of different parameter sets. The large number of parameters of this model makes the use of the exploration tool relevant.

**Fig 15.**
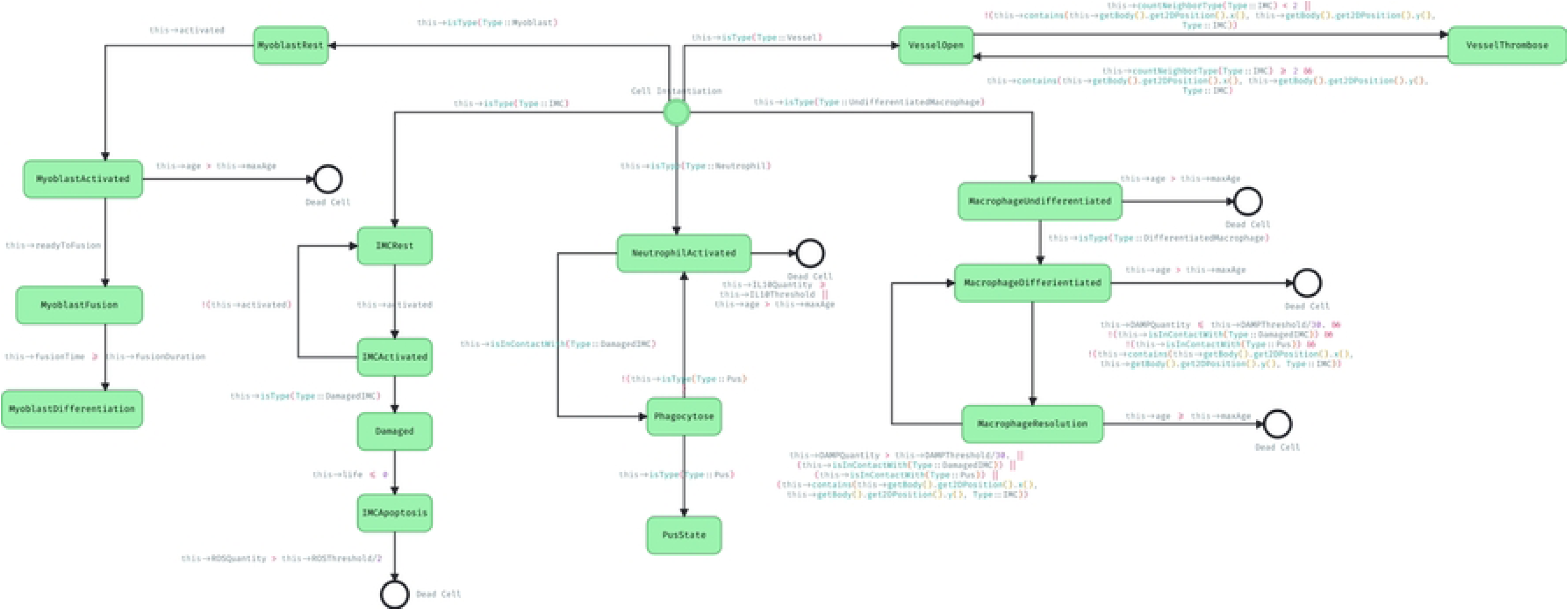
Muscle wound state transition diagram. This diagram reflects the complexity of this model which makes it more easily understandable and thus more easily reproducible.

## Discussion / Conclusion

Agent-based models offer the possibility to study cellular biology at the individual cell scale to explore the basic behavior of cells which are responsible of the emergence of more complex phenomena at the tissue scale. To facilitate and improve the interaction and intercomprehension between modelers and biologists, we proposed a participatory methodology based on software engineering diagrams. In parallel, we also developed a web platform to support and apply our methodology more efficiently. The platform and the methodology were both tested with the implementation of models from the literature and from workshops with biologists. The code of the platform is available here: https://gitlab.com/CogoniXkill/isicell.

Compared to reproducing models from the literature, as computer scientists, developing models with the enlightenment of an expert is far easier. Interacting with biologists provides an expert point of contact allowing for direct explanation of implicit phenomena and characteristics of the modeled system. Integrating biologists in all stages of prototyping through workshops allows for a better communication and intercomprehension with the modelers, thus preventing from over- and misinterpretations. The knowledge of the biologists enables to clarify aspects that were not explicit for the modelers and to qualitatively evaluate the credibility of the produced model. Their expertise is the key to build relevant models in shorter time. Focusing on the behavior also helps the interactions with the biologists.

The platform is part of the agent-based paradigm and thus avoids the possibility of using another type of model. However, the difficulty to evaluate and quantify the efficiency and relevance of a methodology compared to another makes nearly impossible to prove that a paradigm is better than another for a specific model [48]. Nevertheless, in a goal to facilitate interactions, ABMs offer a great foot ground for a better intercomprehension between biologists and modelers. Moreover, the platform is currently biased by the modelers that are working with it but the opening of the platform to a wide public might resolve this problem by providing more user feedbacks.

Currently, using the ISiCell platform for a modeler requires a small amount of training mainly due to the few C++ code to be written to implement compartmental blocks. AI-based Pair programming such as github copilot [49, 50] would be interesting tools to facilitate the handling of the platform for new modelers especially for the creation of new modules. More generally, the increasingly widespread use of Large Language Models (LLMs) [51, 52] such as OpenAI’s GPT-3 [53, 54] which are able to generate coherent text depending on a natural language request, could lead to the development of a myriad of coding tools making programming even more efficient. These AIs already show interesting results for pair-programming or generating code from prompt [55, 56] and could be in a mid-term future interesting tool to improve the usability of ISiCell Builder for modelers.

Although the platform does not need a lot of coding to obtain a functional model, it still requires a minimum of C++ knowledge. A way to make this platform more accessible and usable for biologist could be to develop a metalanguage in the same way as the gamma platform and its GAML [57] to simplify the coding for non computer scientists.

Expanding its community is one of the future goal of ISiCell. Involving more modelers in this project will lead to a virtuous and continuous improvement of the platform while leading to the development of new models. These models will bring new modules specifically created to answer new problematics. If the community is active enough, it could lead to the development of ISiCell Hub, a web platform to more easily share models and modules. In the same way as ISiCell Builder, ISiCell Viewer could become modular to accept new visualization and analysis tools.

## Acknowledgments

This work is supported by grants from the Occitanie Region and University Toulouse Capitole (OnkoOptim project), from Bristol-Myers-Squibb (no. CA184-575) and from the AI Interdisciplinary Institute ANITI (funded by the French program “Investing for the Future – PIA3” under Grant agreement no. ANR-19-PI3A-0004). The research has received funding from the European Research Council (ERC) under the European Union’s Horizon 2020 Research and Innovation Programme (Grant agreement No. Syn-951329).

## Notes

### Competing Interest Statement

The authors have declared no competing interest.

## References

1. Southern J, Pitt-Francis J, Whiteley J, Stokeley D, Kobashi H, Nobes R, et al. Multi-scale computational modelling in biology and physiology. Progress in biophysics and molecular biology. 2008;96(1-3):60–89.

2. Gautieri A, Buehler MJ. Multi-scale modeling of biomaterials and tissues. In: Materiomics: Multiscale Mechanics of Biological Materials and Structures. Springer; 2013. p. 13–55.

3. Baker QB. Computational modeling to study disease development: Applications to breast cancer and an in vitro model of macular degeneration. Utah State University; 2015.

4. Materi W, Wishart DS. Computational systems biology in drug discovery and development: methods and applications. Drug discovery today. 2007;12(7-8):295–303.

5. Macal CM. Everything you need to know about agent-based modelling and simulation. Journal of Simulation. 2016;10(2):144–156.

6. An G, Mi Q, Dutta-Moscato J, Vodovotz Y. Agent-based models in translational systems biology. Wiley Interdisciplinary Reviews: Systems Biology and Medicine. 2009;1(2):159–171.

7. Gorochowski TE. Agent-based modelling in synthetic biology. Essays in biochemistry. 2016;60(4):325–336.

8. Hellweger FL, Clegg RJ, Clark JR, Plugge CM, Kreft JU. Advancing microbial sciences by individual-based modelling. Nature Reviews Microbiology. 2016;14(7):461.

9. Macklin P, Frieboes HB, Sparks JL, Ghaffarizadeh A, Friedman SH, Juarez EF, et al. Progress towards computational 3-D multicellular systems biology. In: Systems Biology of Tumor Microenvironment. Springer; 2016. p. 225–246.

10. Macklin P. When seeing isn’t believing: How math can guide our interpretation of measurements and experiments. Cell systems. 2017;5(2):92–94.

11. Ozik J, Collier N, Wozniak JM, Macal C, Cockrell C, Friedman SH, et al. High-throughput cancer hypothesis testing with an integrated PhysiCell-EMEWS workflow. BMC bioinformatics. 2018;19(18):483.

12. Basco-Carrera L, Warren A, van Beek E, Jonoski A, Giardino A. Collaborative modelling or participatory modelling? A framework for water resources management. Environmental Modelling & Software. 2017;91:95–110.

13. Hare M, Letcher RA, Jakeman AJ. Participatory modelling in natural resource management: a comparison of four case studies. Integrated Assessment. 2003;4(2):62–72.

14. Castelletti A, Soncini-Sessa R. Bayesian Networks and participatory modelling in water resource management. Environmental Modelling & Software. 2007;22(8):1075–1088.

15. Bousquet F, Le Page C. Multi-agent simulations and ecosystem management: a review. Ecological modelling. 2004;176(3-4):313–332.

16. Thorne BC, Bailey AM, Peirce SM. Combining experiments with multi-cell agent-based modeling to study biological tissue patterning. Briefings in bioinformatics. 2007;8(4):245–257.

17. Barreteau O, Antona M, D’Aquino P, Aubert S, Boissau S, Bousquet F, et al. Our companion modelling approach. Journal of Artificial Societies and Social Simulation. 2003;6(1).

18. Bommel P, Becu N, Le Page C, Bousquet F. Cormas: an agent-based simulation platform for coupling human decisions with computerized dynamics. In: Simulation and gaming in the network society. Springer; 2016. p. 387–410.

19. Kaner S. Facilitator’s guide to participatory decision-making. John Wiley & Sons; 2014.

20. Jacob RJK. A state transition diagram language for visual programming. Computer. 1985;18(08):51–59.

21. Harel D. Statecharts: A visual formalism for complex systems. Science of computer programming. 1987;8(3):231–274.

22. Dumas M, Ter Hofstede AH. UML activity diagrams as a workflow specification language. In: International conference on the unified modeling language. Springer; 2001. p. 76–90.

23. Arora V, Bhatia R, Singh M. Synthesizing test scenarios in uml activity diagram using a bio-inspired approach. Computer Languages, Systems & Structures. 2017;50:1–19.

24. Loh WL. On Latin hypercube sampling. The annals of statistics. 1996;24(5):2058–2080.

25. Helton JC, Davis FJ. Latin hypercube sampling and the propagation of uncertainty in analyses of complex systems. Reliability Engineering & System Safety. 2003;81(1):23–69.

26. Mueller-Klieser W. Multicellular spheroids. Journal of cancer research and clinical oncology. 1987;113(2):101–122.

27. Amereh M, Edwards R, Akbari M, Nadler B. In-silico modeling of tumor spheroid formation and growth. Micromachines. 2021;12(7):749.

28. Mao X, McManaway S, Jaiswal JK, Patel PB, Wilson WR, Hicks KO, et al. An agent-based model for drug-radiation interactions in the tumour microenvironment: Hypoxia-activated prodrug SN30000 in multicellular tumour spheroids. PLoS computational biology. 2018;14(10):e1006469.

29. Schaller G, Meyer-Hermann M. Continuum versus discrete model: a comparison for multicellular tumour spheroids. Philosophical Transactions of the Royal Society A: Mathematical, Physical and Engineering Sciences. 2006;364(1843):1443–1464.

30. Pascalie J, Luga H, Lobjois V, Ducommun B, Duthen Y. A Checkpoint-Orientated Modelling for Cell Cycle Simulation. In: International Conference on Bio-Inspired Models of Network, Information, and Computing Systems. Springer; 2010. p. 40–47.

31. Pascalie J, Lobjois V, Luga H, Ducommun B, Duthen Y. A checkpoint-orientated model to simulate unconstrained proliferation of cells. In: ECAL. Citeseer; 2011. p. 630–637.

32. Laurent J, Frongia C, Cazales M, Mondesert O, Ducommun B, Lobjois V. Multicellular tumor spheroid models to explore cell cycle checkpoints in 3D. BMC cancer. 2013;13(1):1–12.

33. Bernard D, Mondesert O, Gomes A, Duthen Y, Lobjois V, Cussat-Blanc S, et al. A checkpoint-oriented cell cycle simulation model. Cell Cycle. 2019;18(8):795–808.

34. Van Liedekerke P, Buttenschön A, Drasdo D. Off-lattice agent-based models for cell and tumor growth: numerical methods, implementation, and applications. In: Numerical methods and advanced simulation in biomechanics and biological processes. Elsevier; 2018. p. 245–267.

35. Grimes DR, Kelly C, Bloch K, Partridge M. A method for estimating the oxygen consumption rate in multicellular tumour spheroids. Journal of The Royal Society Interface. 2014;11(92):20131124.

36. Benzekry S, Chapuisat G, Ciccolini J, Erlinger A, Hubert F. A new mathematical model for optimizing the combination between antiangiogenic and cytotoxic drugs in oncology. Comptes Rendus Mathematique. 2012;350(1-2):23–28.

37. Swierniak A, Kimmel M, Smieja J. Mathematical modeling as a tool for planning anticancer therapy. European journal of pharmacology. 2009;625(1-3):108–121.

38. Pappalardo F, Pennisi M, Castiglione F, Motta S. Vaccine protocols optimization: in silico experiences. Biotechnology Advances. 2010;28(1):82–93.

39. Khazen R, Müller S, Lafouresse F, Valitutti S, Cussat-Blanc S. Sequential adjustment of cytotoxic T lymphocyte densities improves efficacy in controlling tumor growth. Scientific reports. 2019;9(1):1–11.

40. Van Liedekerke P, Palm M, Jagiella N, Drasdo D. Simulating tissue mechanics with agent-based models: concepts, perspectives and some novel results. Computational particle mechanics. 2015;2(4):401–444.

41. Almet AA, Maini PK, Moulton DE, Byrne HM. Modeling perspectives on the intestinal crypt, a canonical system for growth, mechanics, and remodeling. Current Opinion in Biomedical Engineering. 2020;15:32–39.

42. Buske P, Galle J, Barker N, Aust G, Clevers H, Loeffler M. A comprehensive model of the spatio-temporal stem cell and tissue organisation in the intestinal crypt. PLoS computational biology. 2011;7(1):e1001045.

43. Vodovotz Y, Chow CC, Bartels J, Lagoa C, Prince JM, Levy RM, et al. In silico models of acute inflammation in animals. Shock. 2006;26(3):235–244.

44. Li NY, Verdolini K, Clermont G, Mi Q, Rubinstein EN, Hebda PA, et al. A patient-specific in silico model of inflammation and healing tested in acute vocal fold injury. PloS one. 2008;3(7):e2789.

45. Carbo A, Bassaganya-Riera J, Pedragosa M, Viladomiu M, Marathe M, Eubank S, et al. Predictive computational modeling of the mucosal immune responses during Helicobacter pylori infection. PloS one. 2013;8(9):e73365.

46. Gopalakrishnan V, Kim M, An G. Using an agent-based model to examine the role of dynamic bacterial virulence potential in the pathogenesis of surgical site infection. Advances in wound care. 2013;2(9):510–526.

47. Wilensky U, Rand W. An introduction to agent-based modeling: modeling natural, social, and engineered complex systems with NetLogo. Mit Press; 2015.

48. Voinov A, Jenni K, Gray S, Kolagani N, Glynn PD, Bommel P, et al. Tools and methods in participatory modeling: Selecting the right tool for the job. Environmental Modelling & Software. 2018;109:232–255.

49. Bird C, Ford D, Zimmermann T, Forsgren N, Kalliamvakou E, Lowdermilk T, et al. Taking Flight with Copilot: Early Insights and Opportunities of AI-Powered Pair-Programming Tools. Queue. 2023;20(6):35–57. doi:10.1145/3582083.

50. Imai S. Is GitHub copilot a substitute for human pair-programming? An empirical study. In: Proceedings of the ACM/IEEE 44th International Conference on Software Engineering: Companion Proceedings; 2022. p. 319–321.

51. Shen Y, Heacock L, Elias J, Hentel KD, Reig B, Shih G, et al. ChatGPT and Other Large Language Models Are Double-edged Swords; 2023.

52. Kung TH, Cheatham M, Medenilla A, Sillos C, De Leon L, Elepaño C, et al. Performance of ChatGPT on USMLE: Potential for AI-assisted medical education using large language models. PLOS Digital Health. 2023;2(2):e0000198.

53. Dale R. GPT-3: What’s it good for? Natural Language Engineering. 2021;27(1):113–118.

54. Zhang M, Li J. A commentary of GPT-3 in MIT Technology Review 2021. Fundamental Research. 2021;1(6):831–833.

55. Vaithilingam P, Zhang T, Glassman EL. Expectation vs. experience: Evaluating the usability of code generation tools powered by large language models. In: Chi conference on human factors in computing systems extended abstracts; 2022. p. 1–7.

56. Xu FF, Alon U, Neubig G, Hellendoorn VJ. A systematic evaluation of large language models of code. In: Proceedings of the 6th ACM SIGPLAN International Symposium on Machine Programming; 2022. p. 1–10.

57. Taillandier P, Gaudou B, Grignard A, Huynh QN, Marilleau N, Caillou P, et al. Building, composing and experimenting complex spatial models with the GAMA platform. GeoInformatica. 2019;23:299–322.

